# « Idol with feet of clay »: reliable predictions of forest ecosystem functioning require flawless climate forcings

**DOI:** 10.1101/2021.03.02.433613

**Authors:** M. Jourdan, C. François, N. Delpierre, N. Martin St-Paul, E. Dufrêne

**Affiliations:** Université Paris-Saclay, CNRS, AgroParisTech, Ecologie Systématique et Evolution, 91405, Orsay, France; Institut Universitaire de France (IUF); Ecologie des Forêts Méditerranéennes (URFM), INRAE, F-84914, Avignon, France

**Keywords:** Process-based models, climate change, regional scale, photosynthesis, respiration

## Abstract

Climate change affects various aspects of the functioning of ecosystem, especially photosynthesis, respiration and carbon storage. We need accurate modelling approaches (*impact models*) to simulate the functioning, vitality and provision of ecosystem services of forests in a warmer world. These impact models require climate data as forcings, which are often produced by climate models comparing more or less well with observational climate data. The bias percentage of the climate forcings propagates throughout the modeling chain from the climate model to the impact model.

In this study, we aimed to quantify these bias percentage, addressing three questions: (1) Do the impact model predictions vary when forcing it with different climate models, and how do the predictions under climate model vs. observational climate forcing differ? (2) Does the variability in the impact climate simulations caused by climate forcings fade out at large spatial scale? (3) How the fact of using simulated climatic data affects the process-based model predictions in the case of stressful events?

To answer these questions, we present results obtained over the historical period (e.g. 1970-2010) with the CASTANEA ecophysiological forest model and use the data from three climate models. Our analysis focuses on French forests, studying European beech (*Fagus sylvatica*), temperate deciduous oaks (*Quercus robur* and *Q. petraea*), Scots pine (*Pinus sylvestris*) and spruce (*Picea abies*) monospecific stands.

We show that prediction of photosynthesis, respiration and wood growth highly depends on the climate model used, whether debiased or not, and also on species and region considered. Overall, we observed an improvement of prediction after a monthly mean bias or monthly quantile mapping correction for three model considered, but not with the same success. Then we highlighted a large variability in the processes simulated by the impact model under different climate forcings when considering the plot (i.e. scale of a few hectares) scale. This variability fades out at larger scale (e.g. the scale of an ecological region, i.e. 100 km^2^), owing to an aggregation effect. Moreover, process predictions obtained under different climate forcings are more variable during driest years. These results highlight the necessity to quantify bias and uncertainties in climate forcings before predicting fluxes dynamics with process-based model.

## 1 Introduction

In the Northern hemisphere, climate change will lead to increased temperature and spatially heterogeneous changes in precipitation regime as well as more frequent and intense extreme climatic events in the forthcoming decades (Pachauri et al., 2014), particularly with strong drought and/or thermal stresses. Such changes, and notably extreme drought events (Tramblay et al., 2020), may be very damaging for European ecosystems functioning (Maracchi et al., 2005) and especially for forests. Warmer and drier conditions can lead to medium or long-term damage (Bréda et al., 2006; Linares et al., 2010). They impact fundamental biological processes involved in the energy, carbon and water cycles, (Anderegg et al., 2012; Dusenge et al., 2019; van der Molen et al., 2011), and may lead to massive dieback, as already reported (Ogaya et al., 2020; Senf et al., 2020). Moreover stressful climatic events can also increase the vulnerability of forest stands to pathogen attacks (Desprez-Loustau et al., 2006) or fire risks (Dale et al., 2001).

It has become critical to take an interest in the forest functioning changes in the coming decades, in order to maintain forest cover and ecosystems services over the long term. It is notably important to quantify forest productivity at scales ranging from the plot (sylvicultural unit) to regional scales (wood supply basin) taking into account the climate. This would help in decision-making and quantification of their contribution to carbon neutrality. Many studies have already addressed climate change effects on the fundamental processes governing forest ecosystems (Lindner et al., 2010; Sperry et al., 2019), describing positive effects (Gunderson et al., 2009) and in more extreme conditions negative effects (Sperry et al., 2019). Most studies used data from the last decades (second half part of the last century) (Zimmermann et al., 2015) or spatial stress gradients (latitudinal, Frenne et al., 2013; Pretzsch et al., 2010, or altitudinal, Marcora et al., 2017) to represent processes evolution during stressful climatic conditions intensification. These approaches is limited in terms of studied ecosystems: not all forests types can be studied on a wide climatic gradient. Precise and extensive climatic datasets are not very old (only a few decades long, for example French mesoscale SAFRAN analysis system (Durand et al., 1993) in France is available only from 1959, for example), and spatial gradients are limited to existing conditions. Experimentation allow also studying global change effect on forest physiological processes, such as CO2 fertilizing effect on photosynthesis (Walker et al., 2020) or increasing temperature effect on tree bud burst (Fu et al., 2012; Wang et al., 2019). These approaches are very useful, but remain rare because such experiments are difficult to implement and manage and/or limited to young trees.

Simulation models are a promising tool and their contribution are crucial: it makes it possible to go beyond the intrinsic limits of field studies. Modeling in future climate has already fueled the scientific debate (García‐Valdés et al., 2020) and provided keys for forest managers practices (Gupta & Sharma, 2019; Lindner et al., 2000). In the large majority of studies this type of forecasting is carried out in the contemporary period with empirical growth models (BICAFF project, Valade et al., 2018 or MARGOT model, Wernsdörfer et al., 2012). These models are relevant in constant climate conditions, but they are not able (by construction) to take into account the influence of climate change on the processes of forest functioning (passing over the fertilizing CO2 effect on photosynthesis, Walker et al 2020, or increasing temperature effect on photosynthesis and respiration, Dusenge et al., 2019, already show in literature), especially their productivity (Boisvenue & Running, 2006; Reyer et al., 2014), but also photosynthesis and respiration.

Process-based models could induce many advances in forest functioning predictions. Indeed they allow to integrate the direct effects of the climate (temperature, precipitation, radiation or relative humidity) on the processes involved in the functioning of ecosystems (photosynthesis and respiration for example) and therefore make it possible to project their functioning under changing conditions, not yet observable on the field. It is not known whether their outputs are significantly affected by the precision of the climate forcings required to run them. For example, Palma et al. (2018) suggests little process uncertainties due to climate data used (with BiomeBGC model), while Stéfanon et al. (2015) and (Glotter et al., 2014) suggests conversely a strong effect of uncertainties due to climate input. Thus it is important to understand uncertainty level associated with the climate model used. However, uncertainties is accumulated along a “cascade of uncertainty” (Lindner et al., 2014; Reyer, 2013), composed of uncertainty at each step of the chain, in our study. In a first step, assumptions about the development pathways of future societies was developed in IPCC report (Pachauri et al., 2015). These storylines allowed developing RCP scenarios (Moss et al., 2010; van Vuuren et al., 2011). The projected greenhouse gas emissions then drive GCMs to provide global climate change scenarios, which are downscaled to lower resolutions using RCMs (called uncorrected modelling climate in our study). To use that data at the forest stand level, a further downscaling to forest stands scale is required (at 8 km in our corrected modelling climate data). Then the data may be used as input into a stand-level forest model. In our work, we try to quantify the uncertainty due to climatic input data, i.e. this step of the “cascade of uncertainty”. In fact global and regional climate model (GCM and RCM) outputs are often used as climate forcing for ecological impact models, and this potentially results in large cumulative errors because information and error are passed sequentially along the modeling chain from GCM to RCM to impact model.

This is especially important when moving from micro-local (plot) to regional scales (environnemental and sylvicultural unit). Indeed the change of scale involves other factors which vary (age, total biomass, soil fertility) that can compensate the errors and thus limit the contribution of the climate in the predictability of process. Moreover, it is crucial to evaluate and decrease bias percentage of process estimation (*GPP*, respiration or *AWBI,* aboveground wood biomass increment) during stressful year, in order to correctly predict furthers changes: (1) because extreme stressful conditions are predicted to be more frequent in next decades (Polade et al., 2017), (2) climate models have more difficulty reproducing extreme climatic conditions (Iizumi et al., 2017) and impact models have difficulties to estimate their effects (Albrich et al., 2020), and (3) ecosystem responses can vary greatly when a threshold is exceeded (Dorman et al., 2015; Meir et al., 2015), which is a situation that is most often encountered in stressful years. There are at least two steps to this assessment work: estimating error induced by climate data quality used as input and estimating stress response quality in the impact model (van Horssen et al 2002). Here we will only focus on the first part.

Our objective in this study is to evaluate the ability of different climatic simulation models (as opposed to observational data) to be used as input data for process-based model simulations. We propose here to evaluate process-based model simulation on the historical period (1960-2010), to estimate the likely error of predictions, with different modelling climatic data. Palma et al. (2018) and Stéfanon et al. (2015) conducted earlier similar studies. Contrary to these studies, we worked on climate model use with annual (or pluriannual) fluxes comparison between simulations, using historical climatic data, and modelling climate simulations, as opposed to forest distribution or forest stock considered in these previous works. Understanding bias allow improving predictions. Then forest process based modelling applications would be multiple in prediction in climate change context: deepen the knowledge of process variations, estimate change in ecosystems services (as productivity or carbon storage) or test different forest management (e.g. Garcia-Gonzalo et al., 2014; Palma et al., 2015) to promote the most resilient. In this study, we answered to: (1) Do the impact model predictions vary when forcing it with different climate models, and how do the predictions under climate model vs. observational climate forcing differ? (2) Does the variability in the impact climate simulations caused by climate forcings fade out at large spatial scale? (3) How the fact of using simulated climatic data affects the process-based model predictions in the case of stressful events?

Our analysis focuses on French forests, studying European beech (*Fagus sylvatica L.*), temperate deciduous oaks (*Quercus robur* and *Q. petraea*), Scots pine (*Pinus sylvestris*) and spruce (*Picea abies*) monospecific stands. These dominant and representative stand of temperate forests in Europe and in France (50% of French wood volume according IFN, French Forest National Inventory, https://inventaire-forestier.ign.fr/) are important for timber or patrimonial reasons. To treat this topic, we used the process-based model CASTANEA (Dufrêne et al., 2005) to test relevance of 3 different climatic models (including correction or not, i.e. six climate cases) on past period, comparing with historical climatic data (French mesoscale SAFRAN analysis system, Durand et al., 1993). Comparison of CASTANEA simulations using different climate models (with and without bias corrections) as input allow to quantify percentage of bias associated to different modelling climate data and evaluate their relevance in prediction with process-based models. Comparing corrected and uncorrected modeling data effect on CASTANEA prediction allows to assess the importance of correction.

## 2 Materiel and Method

### 2.1 Process-based model description

The process-based model CASTANEA (Dufrêne et al., 2005) aims at simulating carbon and water fluxes and stocks of even-aged regular monospecific forest stand at the rotation timescale (several decades). In this study, CASTANEA was used to simulate the annual stand photosynthesis, respiration and wood growth of monospecific stand (as Delpierre et al., 2012; Guillemot et al., 2014b), but others studies focused on allocation (Davi et al., 2009a; Guillemot et al., 2017) and also on water fluxes (Davi et al., 2006).

In brief, stand simulated by CASTANEA comprises four functional compartments: foliage, woody biomass (including stem, branches and coarse roots), fine roots and a pool of carbohydrate reserves. The canopy is considered homogeneous horizontally and vertically sub-divided into a given number of layers (i.e. about 30), each of them enclosing a constant amount of leaf area. The model performs well in simulating the interannual fluctuations of CO2 forest-atmosphere exchanges (Delpierre et al., 2012) and the spatial and interannual variations of wood growth (Guillemot et al., 2014b). A complete description of CASTANEA is given in Dufrêne et al. (2005), with subsequent modifications from Davi et al. (2009), Delpierre et al. (2012) and Guillemot et al. (2014, 2017). Carbon allocation to wood growth is determined daily as a fraction of net primary productivity (*NPP*) using allocation coefficients.

Allocation coefficients are related to stand age and species, as well as to the current and previous year water stress (Guillemot et al., 2014). The module of C allocation to wood growth was preliminarily calibrated and validated at plot scale on RENECOFOR (French National Network monitoring Forest Ecosystems) tree ring series (over the 1970-1990 period, see Guillemot et al., 2014b) and IFN (French National Inventory) inventory (see Appendix S1).

### 2.2 Input data

#### 2.2.1 Studied species

We considered five species in this study: common beech (*Fagus sylvatica*), Norway spruce (*Picea abies*), pedunculate and sessile oaks (*Quercus robur* and *petraea*) and Scots pine (*Pinus sylvestris*). These species are widespread in the French territory, but they are also major species in European temperate forest (San-Miguel-Ayanz et al., 2016), and widely studied. They represent an important percentage of forest volume (IFN, *Le Memento*, 2019): 10% for beech, 11% for pedunculate oak, 11% for sessile oak, 5% for Scots Pine and 8% for spruce (i.e. 45% of total volume of French forest). It also represent economic issues with 18,4 ± 2,9 Mm3/an between 2009-2017 (IFN, *Le Memento*, 2019).

Beech and oaks are broadleaved species while spruce and Scots pine are coniferous species. Beech and spruce are late-successional and shade tolerant species. Spruce is very sensitive to high temperatures in summer. Beech and sessile and pedunculate oaks (Lobo et al., 2018) are sensitive to water stress, when Scots pine tolerate drier conditions (Pasta et al., 2016) and are more early succession and light-demanding species. These five species allowed studying various types of forest, thanks to physiological and ecological differences between species.

In the model, pedunculate and sessile oaks have the same parameterization, but they were distinguished in this study because their spatial distribution and local stations in France are different.

#### 2.2.2 Climate data

CASTANEA model works at an hourly time step. It can be forced with daily climatic data that are internally rescaled at an hourly time step. The daily data used here were radiation (*Rg*), precipitation (*Prec*), daily mean, minimum and maximum of temperature (respectively *T, Tmin, Tmax*) and relative humidity (*RH*) and mean wind speed (*W*). We used two types of data. First a reference dataset which comes from the French mesoscale SAFRAN analysis system (Système d’Analyse Fournissant des Renseignements Atmosphériques à la Neige or in English System of Analysis Providing Atmospheric Information to Snow, Vidal et al., 2010). Secondly, we also used modelled climatic series extracted from Eurocordex data MPI (GCM MPI-ESM-LR downscaled with RCM RCA4), Hadgem (GCM MOHC-HadGEM2-ES downscaled with RCM RCA4) and CNRM (GCM CNRM-CERFACS-CNRMCM5 downscaled with RCM RCA4) models data (see two next paragraphs and Appendix S8), over whole Metropolitan France for the 1961-2010 period.

##### Historical climate data: SAFRAN

The SAFRAN product is a mesoscale analysis system, developed by Météo-France, which reconstructs vertical atmospheric profiles on regular horizontal 8km grid. It spatializes large-scale general circulation models data, combined with observed data. SAFRAN first analyzes temperature, wind, air humidity, incident solar radiation and cloudiness (Quintana-Seguí et al., 2008). For each geographic unit, a draft model is compared to the observations in order to verify the consistency of the observations. The analysis is then performed using valid observations (Vidal et al., 2010). SAFRAN analyses therefore have the advantage of providing at high spatial resolution all the climatic variables necessary for the simulation with CASTANEA over the period 1960-2010, and can serve as climatic reference in our study. SAFRAN has already been validated over French metropolitan territory produces unbiased estimates of precipitation, solar radiation, speed wind, relative humidity and temperature (Quintana-Seguí et al., 2008).

##### Modelled climate data

We decided to work on daily climate data produced in the frame of the EURO-CORDEX initiative (Kotlarski et al., 2014) and processed by Fargeon et al. (2020). CORDEX simulations result from a dynamical downscaling by the coupling of Global Climate Models (GCMs) with Regional Climate Models (RCMs). The selection of the GCM-RCM couples we used was based on: availability in the archiving system of CORDEX simulations (European System Grid Federation, ESGF), antecedent model validation for the area of interest, maximization of the expected differences between models while remaining in the plausible zone.

Finally, we worked with three GCM models (MPI-ESM-LR, MOHC-HadGEM2-ES, CNRM-CERFACS-CNRMCM5) models data downscaled with one RCM model (RCA4) at 50 km (De Cáceres et al., 2018; Fargeon et al., 2020; McSweeney et al., 2015). We used only one RC since these models have a marginal effect compared to GCM on modelling climate data (Glotter et al., 2014). The three selected runs, as well as a description of their projected climatic changes, are summarized in Appendix S8.

To summarize Fargeon et al. (2020) identify a significant bias of uncorrected climatic simulations over the past period (1995–2015) for meteorological variables of interest (*T*, *RH*, *Rg*, *Prec* and *W*) for our study, compared to SAFRAN (Vidal et al., 2010). Hence, projections based on biased data might misestimate anomalies and scenario impact. Consequently, a statistical downscaling and bias corrections were performed, using the SAFRAN reanalysis as reference observational data (8 km resolution), using standard methods based on monthly mean bias or monthly quantile mapping (Bedia et al., 2014; Ruffault et al., 2014), which strongly reduced the bias in model outputs. Temperature and radiation were corrected using a mean bias and precipitation, wind speed and relative humidity were corrected using quantile mapping.

In this study, we wanted (objective n°1, see above) to compare CASTANEA simulations using six different climate data sets (MPI, CNRM and Hadgem, with and without corrections), in order to evaluate the outputs of process-based model CASTANEA with different climatic modelling data.

#### 2.2.3 Inventory

To initialize state variables and stand characteristics of the CASTANEA model, we used a sub-dataset of French National Forest Inventory. Because CASTANEA modelled only monospecific, regular and even-aged stands, we considered only stands constituted of more than 70% of objective species (in basal area and density). In addition to have regular inventories, we kept only trees of the dominant stratum in plot of IFN inventory. Plots stand younger than 20 years are excluded (less than 8% of monospecific plots previously selected), because CASTANEA did not manage the youngest stand. We used inventory data (over 2005-2017, available at https://inventaire-forestier.ign.fr/)

In the final dataset we have 2379 plots of pedunculate oak, 2178 plots of sessile oak, 2196 plots of beech, 1364 plots of spruce and 1834 plots of Scots pine.

Then two regional scales SER (meso-regional scale, “Sylvo-Eco-Region”, i.e. region with forest and ecological coherence, Cavaignac 2009) and GRECO (macro-regional scale, Bonheme 2021) will be used to aggregate results at plot scale.

### 2.3 Simulation design and process studied

CASTANEA simulations are done over the period 1960-2010, initialized with the inventories previously detailed and the SWHC (soil water holding capacity) from Q-div project (Badeau, 2011). Each species is treated separately. It is necessary to minimize because we only want to look at the effect of the climate and not other evolving factors. Stressful years will not occur at the same time for SAFRAN and for climate modeling data, while trees would response differently depending on their age and biomass, therefore age and biomass are kept constant, per decade (e.i. 1960-1970, 1970-1980, 1980-1990, 2000-2010) with a reinitialization of stand age and biomass. This duration makes it possible to have a minimal change in age and biomass, and to be able to take into account the lag-effects of climate.

We focused on several output variables of CASTANEA, to evaluate the use of climate models described earlier (see section 2.2.2), characterizing ecosystems functioning and being the added value of process-based model: photosynthesis (*GPP*), maintenance respiration (*Resp*) allowing to study carbon cycle, aboveground wood biomass increment (*AWBI*), which interest managers (in the context of an applied use of climate models).

### 2.4 Analyses

#### 2.4.1 CASTANEA calibration on French territory

First, we evaluated CASTANEA on French territory, thanks comparison between simulated annual radial increment (*RI_sim_*) and annual radial increment of IFN inventory, between 2006-2017 (*RI_obs_*).

The evaluation was performed for each species comparing *RI_sim_* and *RI_obs_*, for each SER, averaging plot value, in order to decrease relative weight of outlier structure (as very low density or age, or extrem SWHC). We also compared *AWBI_sim_* and *AWBI_obs_* (see Fig 1), using radial increment to *AWBI* conversion developed in (Guillemot et al., 2014a). Total observed volume is computed thanks observed radial increment, diameter and allometries used in CASTANEA: relation between age and dominant height -from Bontemps et al., (2007) for beech stand, Bontemps et al., (2012) for oaks, Seynave et al., (2005) for spruce and Palahı́ et al., (2004) for Scots pine-, relation diameter, height and volume (Vallet et al., 2006) among in other. With this volume and the wood density used in the model (660 for oaks, 550 for beech, 379 for spruce and 440 for Scots pine) we can compute *AWBI,_obs_* and compare it with *AWBI,_sim_.*

**Figure 1.**
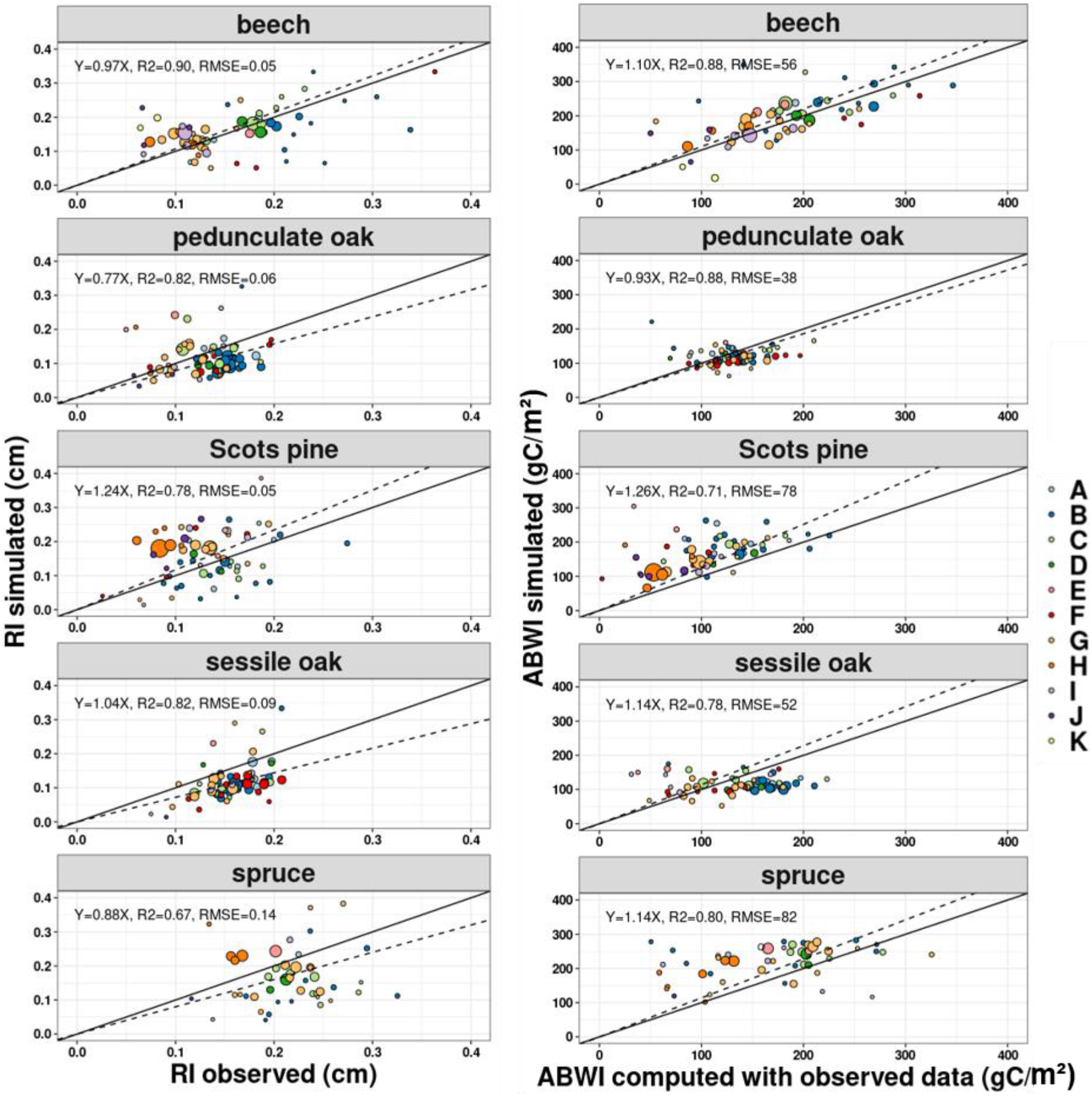
Graph represents annual radial increment in cm (left) and NPP_wood_ in gC/m^2^ (right) simulated vs observed, by species. Each point represents mean plot value for each SER (see Figure S1). Size represents number of plot by SER and colour indicates GRECO. Dashed line represent linear model Var_sim_~ Var_obs_ and regression equation is indicated with R-squared for each species (coefficients are always significant, pvalue<0.0005). The black line is the line 1:1.

#### 2.4.2 Climate models vs. SAFRAN: bias of impact model outputs for non-extreme years

##### Plot scales

We wanted to compare impact model outputs (*AWBI*, *GPP*, *Resp*) resulting from simulations with climate models (*mod*) (corrected: *MPI_corrected_, CNRM_corrected_, Hadgem_corrected_*, or uncorrected: *MPI_raw_ CNRM_raw_ Hadgem_raw_*) and impact model outputs from simulations with SAFRAN data (*saf*), for each species separately. For each plot, impact model outputs were averaged over 1960-2010 to allow statistical comparison between climate models and SAFRAN data.

To quantify percentage of predictions bias, we computed bias between simulations with climatic model and simulations with SAFRAN, by variable (*V*), model (*mod*) and species (*sp*), for each plot *i*:

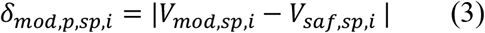

We compared absolute variable bias when simulation were done with uncorrected climatic simulated data (*δ_raw_*) or with corrected climatic simulated data (*δ_corrected_*), plotting *δ_corrected_* in function of *δ_raw_*.

Although validation was done at plot scale, we decided to work at SER (Sylvo-Eco Region or Forest Ecological Region) scale for each species (see Appendix S2) in order to reduce statistical effect of marginal stand (with very low or very high density, for example).

We computed bias percentage to quantify percentage of bias using climatic model compared to the reference (simulation with SAFRAN data). We did it for every model (with and without correction):

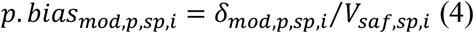

##### Regional Scales

To study scale effect on quality of prediction (objective n°2, see above), we focused on two regional scales, SER and GRECO, to understand aggregation effect on these two scales. At the regional level, stand age distribution could have a strong effect on prediction because processes are strongly affected by stands age in CASTANEA and can interact with climate effects. In the following analyses we considered four age classes (20-60, 60-120, 120-180, >180). For more details see Appendix S3.

We compute the proportion of bias compared to variable level at regional scale as in eq 3, using total surface of the species (*S_sp_*), mean level mean level per hectare of variable V for all plots *i* of a given age class (*mean_i_(V)*) and proportion of this age class (%_sp_,_s_,_age_):

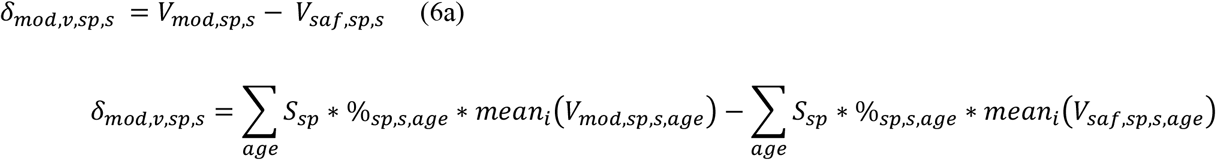

which reduces to:

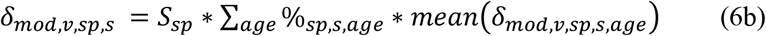

The bias percentage was also calculated:

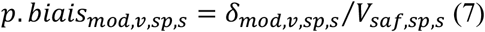

We computed also percentage of bias at a larger scale (GRECO level and France level).

#### 2.4.3 Process bias to SAFRAN simulation for stressful years

For this part (objective n°3, see above), we decided to test the effects of climate forcings on CASTANEA predictions during extreme events. Then we focused on bias estimation on the most stressful years over the period (1960-2010). We selected the driest years of our study period for each dataset, each model and each GRECO (Appendix S11). We use the *SPEI* to identify these years (Vanoni et al., 2016). *SPEI* is derived of the *SPI* (Standardized Precipitation Index, Guttman, 1999), and represents a climatic water balance (Thornthwaite, 1948) calculated at different time scales, using the monthly (or weekly) difference between precipitation and *PET* (Vicente-Serrano et al., 2013; Vicente-Serrano et al., 2010). We calculated annual *SPEI* using *R* package *SPEI* (Vicente-Serrano et al., 2010), between February and July, hereafter identified as *SPEI_07_*, i.e. the growth period (Vanoni et al., 2016, Jourdan et al 2019). It allowed determining the years with driest growth season of a dataset for each climatic model and SAFRAN dataset (Appendix S4 ant Appendix S11).

For this sub-dataset, we also compared *δ_corrected_* and *δ_raw_* (following equation 2) and computed percentage of variables bias simulated with modelling climatic data (following equation 3), as described in 2.4.2.

## 3 Results

### 3.1 CASTANEA evaluation on French territory

The observed annual radial increment range from 0.05 to 0.3 cm with an average of 0,16 cm (± 0,06) for beech, 0,16 cm (± 0,07) and 0,13 cm (± 0,03) for sessile and pedunculated oaks, 0,13 cm (± 0,04) for Scots pine and 0.22 cm (± 0,06) for spruce (Fig 1-a). Values range are equivalent for simulations with: 0,16 (± 0,06) for beech, 0,13 (± 0,14) and 0,12 (± 0,07) for sessile and pedunculate oak, 0,17 (± 0,11) for Scots pine and 0.21 (± 0,15) for spruce.

The annual *AWBI* computed from observed data range from 0 to 400 gC/m^2^/y with an average of 182 (± 64) for beech, 113 (± 28) and 131 (± 29) for sessile oak and pedunculate, 169 (± 25) for Scots pine and 220 (± 50) for spruce (Fig 1-b). Values range are equivalent for simulations with: 196 (± 68) for beech, 131 (± 44) and 119 (± 25) for sessile and pedunculate oak, 107 (± 46) for Scots pine and 171 (± 66) for spruce.

The overall performance of the model is equivalent to what is found in literature (see Palma et al., 2018 or Forrester et al., 2017): the slope of 0.97 (±0.22) obtained in our study is comparable to 1 (±0.17) in Forrester et al (2017). The results vary between species. Results are different for monospecific Scots pine where stands radial increments and annual *AWBI* are slightly overestimated by the model (respectively with slope of 1.14 and 1.4). This observed inaccuracy in Scots pine simulations may be explained by the fact that Scots pine carbon fluxes were calibrated with fluxes site data from the northern part of the distribution area (Hyytiälä, in Finland; Delpierre et al., 2012) while wood growth were calibrated with data from the southern part of distribution area (French national Inventory, IFN). To a lesser extent, annual radial increments and annual *AWBI* fluxes of pedunculate and sessile oaks are slightly underestimated (respectively with slope of 0.74 and 0.70 for annual radial increments and with slope of 0.87 and 0.79 for annual *AWBI* fluxes).

### 3.2 Climate models vs SAFRAN: bias of impact model ouputs

Photosynthesis simulated is in average 1500 gC/m^2^/year for oaks, 1250 gC/m^2^/year for beech, 1750 gC/m^2^/year for Scots pine and 2250 gC/m^2^/year for Spruce (Fig. 2). Photosynthesis simulated with Hadgem climatic model without correction seems lower than others simulations, for every species (except for Spruce).

**Figure 2.**
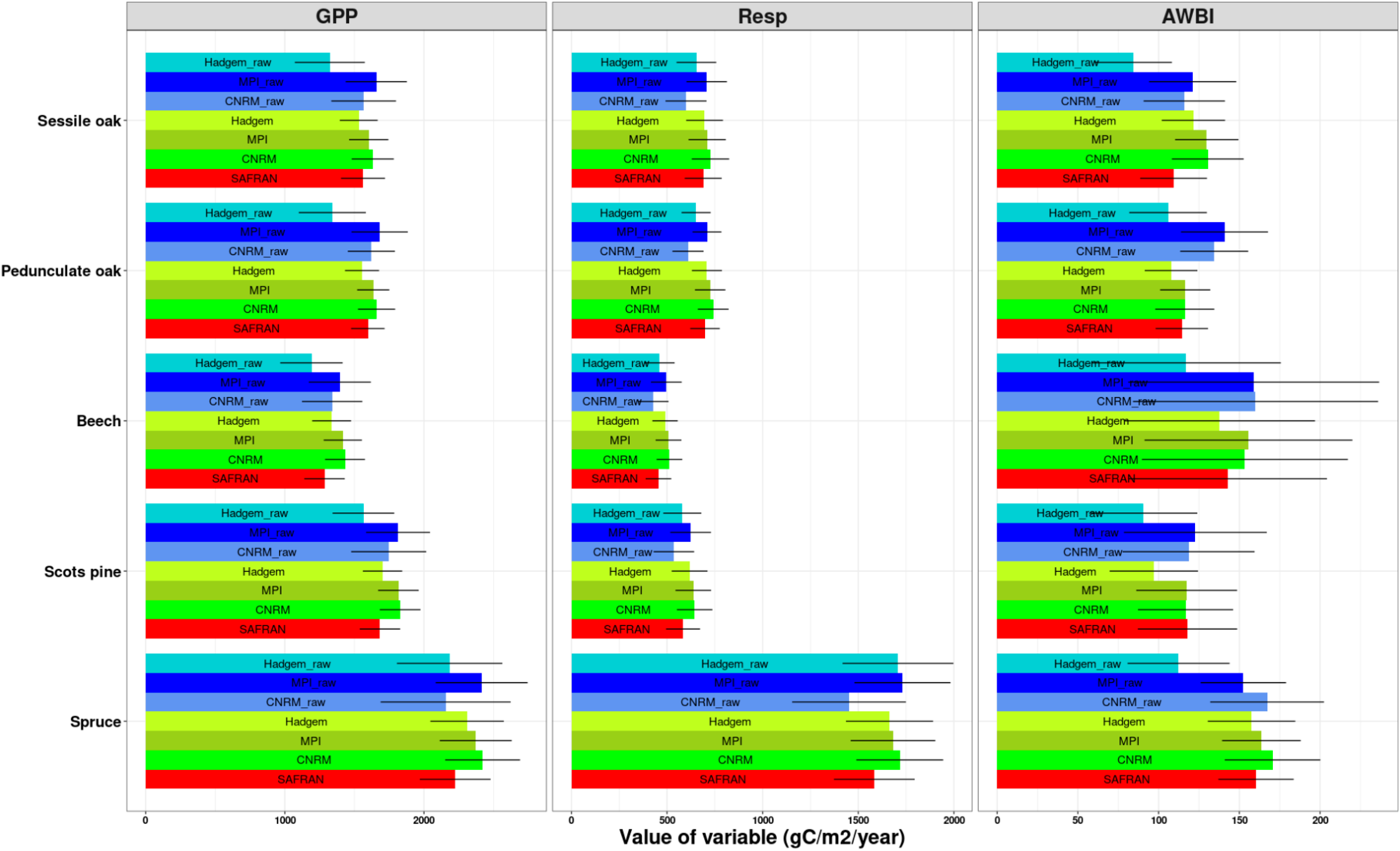
Mean values of NPP_wood_, GPP and Respiration for period 1960-2010, by species and origin of data used: simulation with SAFRAN (red), model before correction (blue) and model after correction (green). Origin of climate data are also written on barplot.

Respiration simulated is in average 700 gC/m^2^/year for oaks, 500 gC/m^2^/year for beech, 600 gC/m^2^/year for Scots pine and 1500 gC/m^2^/year for Spruce. Respiration simulated with ^*CNRM*^ climatic model without correction seems lower than others simulations, except for Spruce (Fig. 2).

Wood carbon simulated is in average 120 gC/m^2^/year for oaks, 150 gC/m^2^/year for beech, 120 gC/m^2^/year for Scots pine and 160 gC/m^2^/year for Spruce. Wood carbon simulated with Hadgem climatic model without correction seems lower than others simulations, except for pedunculate oak (Fig. 2).

#### Bias before vs after correction

Processes biases decrease overall when the climatic data are corrected (in average 17 gC/m^2^/y for *AWBI*, 113 gC/m^2^/y for *GPP* and 40 gC/m^2^/y for respiration) and reach an error of 8% in average at plot level. The absolute value of the bias between simulation with SAFRAN and with uncorrected model is greater than the absolute bias with corrected models, for MPI and Hadgem models (respectively 84% and 85% in average any process combine), meaning that climatic corrections decrease the bias to SAFRAN simulations (Fig. 3). This result is more nuanced for CNRM model (only 58% of *SER* shows a bias improvement for photosynthesis and 60% for respiration).

**Figure 3.**
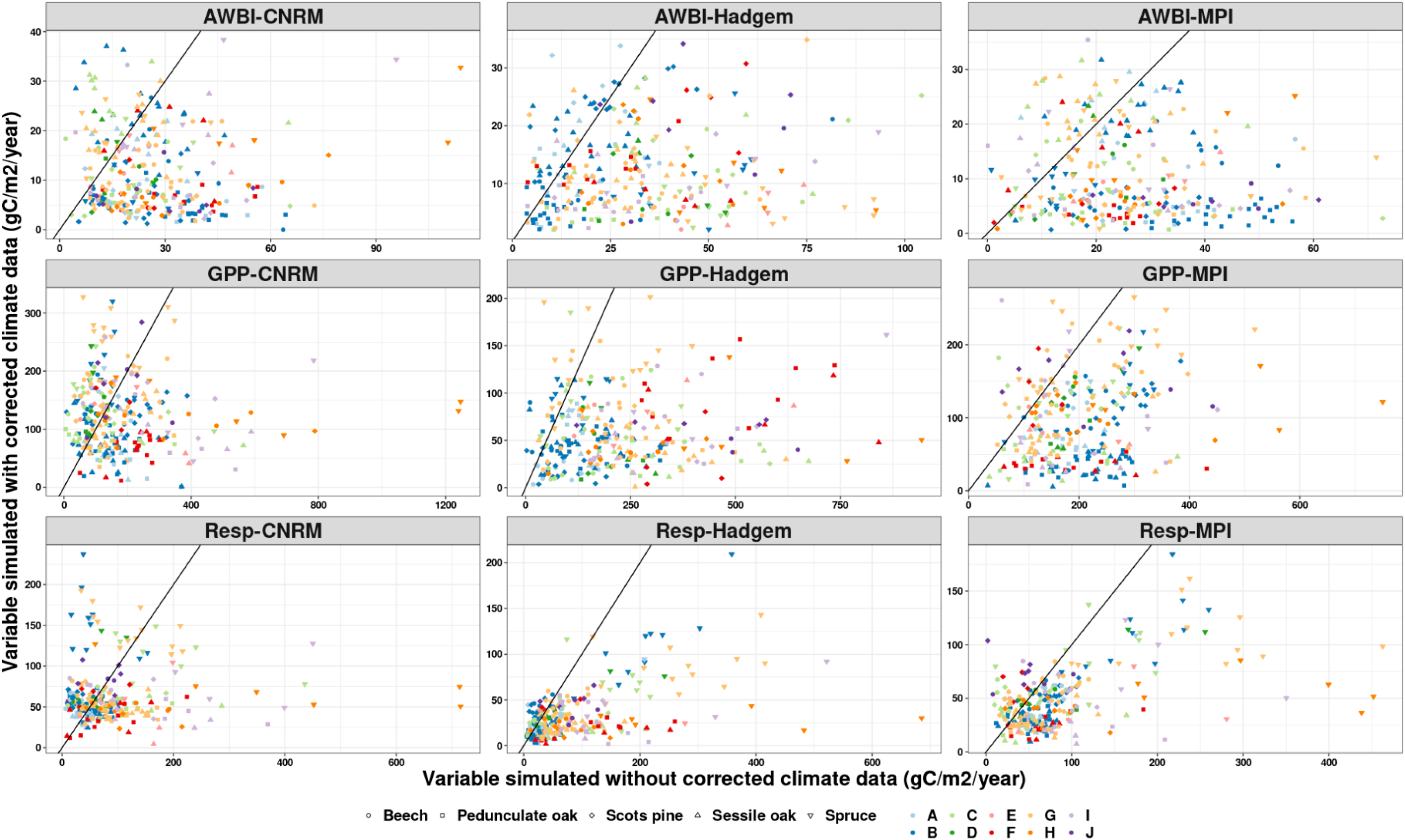
Comparison of process bias between simulation with uncorrected data and corrected data, for each process and each model. Shapes represent species and GRECO are indicated by color. The black line is the line 1:1. Points below the line indicate a decreasing bias between uncorrected and corrected climate data. Points above the line indicate an increasing bias between uncorrected and corrected climate data.

In some few cases, the correction of climate bias did not improve the impact model simulations. Then in these cases the impact model simulation appeared less accurate after climate de-biasing. For the CNRM model, the climate bias correction increases for beech stand (*GPP*, *Resp*), spruce stand (*GPP*), Scots pine stand (*Resp*) and sessile oaks stand (*AWBI*).

#### Does percentage of bias depends on aggregation levels?

Correction of modelling climate data divides by more than two the percentage of variable bias induced by using raw modelling climate data, for each scale and each process (Appendix S12). Percentage of variable bias simulated with corrected modelling climate data differs between scales (plot, SER, GRECO and France), process studied (Fig 4a) and species. At plot level, percentages of variable bias are in average 6% for *GPP*, 7% for *Resp* and 11% for *AWBI* (Appendix S12), but can reach 90% for *AWBI* of monospecific oaks stand and *Resp* and *GPP* of monospecific pedunculate oak and beech stands. At regional scale (SER, GRECO and France), percentages of variable bias are quite the same in average 6% for *GPP*, 7% for *Resp* and 10% for *AWBI* for SER (with a weaker standard error), 6% for *GPP*, 7% for *Resp* and 9% for *AWBI* for GRECO and France (with a weaker standard error). Difference of percentages of variable bias between scales varies depending on monospecific stands: from the highest percentages of variable bias for sessile oak stand *AWBI* (from 11% for plot scales to 9% for France scales) to lowest percentages of variable bias for pedunculated oak stand *GPP* (from 4% for plot scales to 3% for France scales). Percentage of *AWBI bias* are higher than percentage of *GPP* and *Resp* bias, at every scale.

**Figure 4.**
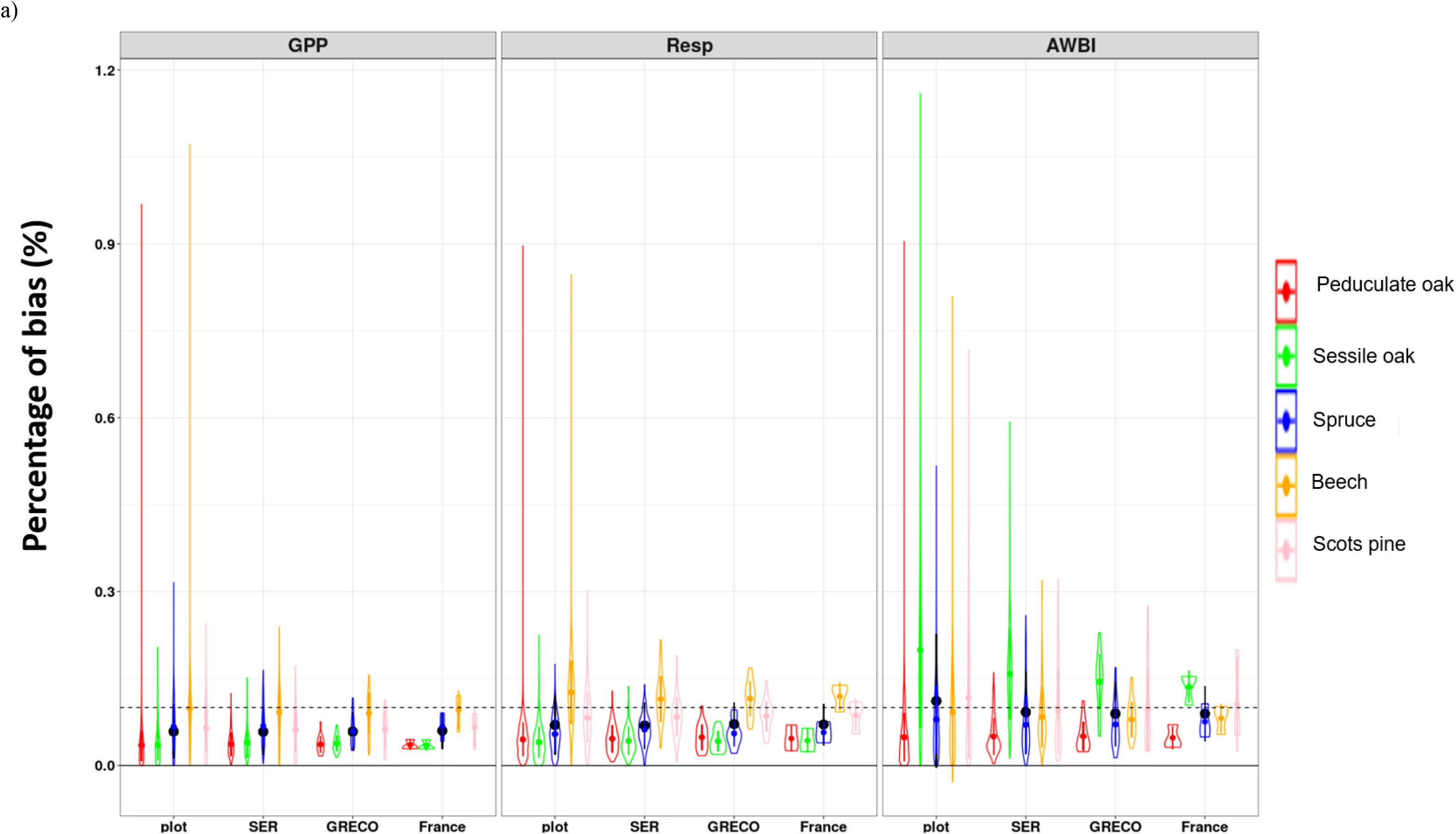
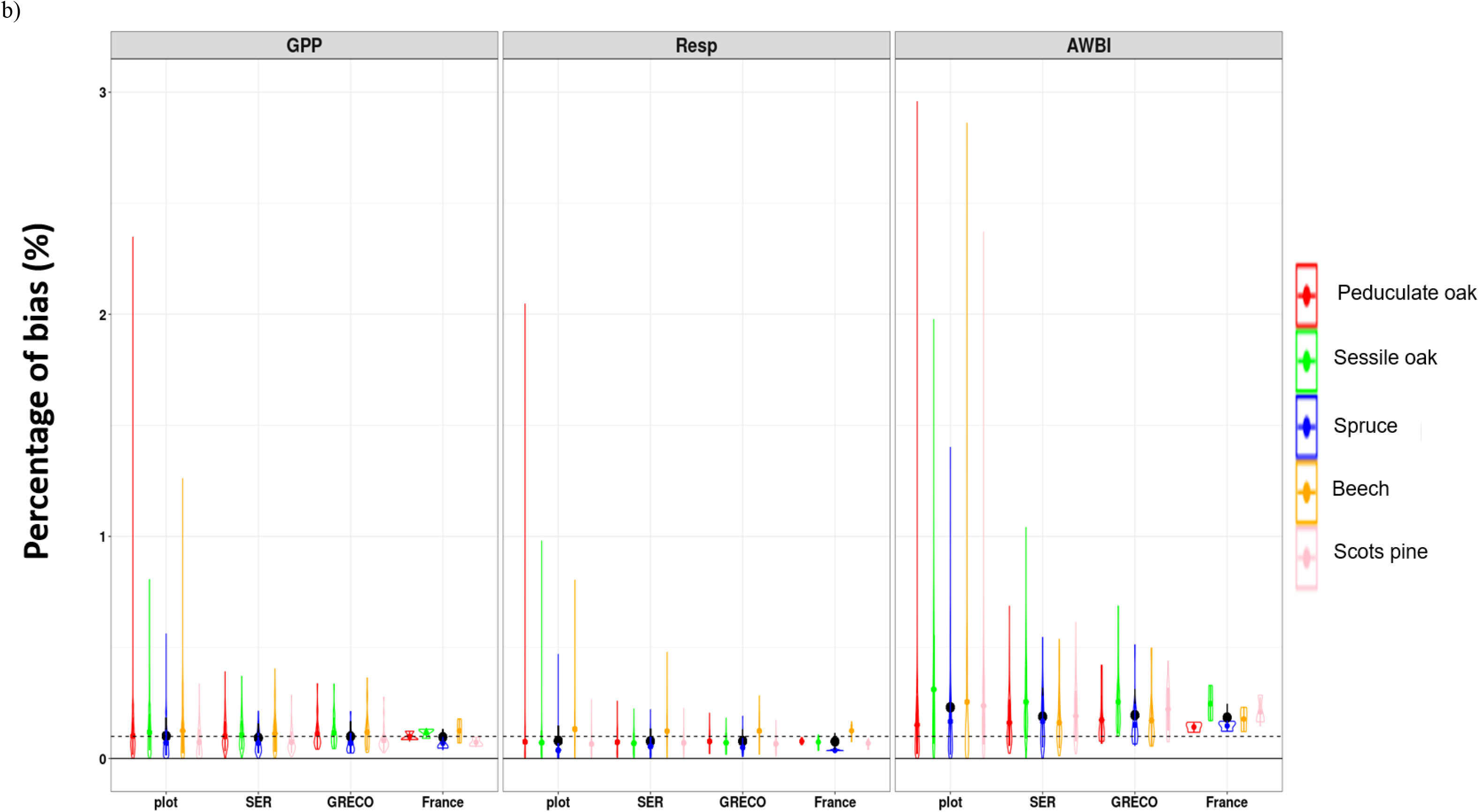
Bias percentage of simulation by model (i.e. percent of bias compared to process levels simulated with SAFRAN), by process (column), scale (x-axis). Black points represents the mean uncertainty by scale and color points represents the mean uncertainty by species and scales. Dashed line represents 10% threshold. Bias percentage of simulation by model (i.e. percent of bias compare to process levels simulated with SAFRAN), by process (column), scale (x-axis) for four driest year in period 1960-2010.). Black points represents the mean uncertainty by scale and color points represents the mean uncertainty by species and scales. Dashed line represents 10% threshold.

#### 3.3 Process bias to SAFRAN simulation for stressful years

##### Bias before vs after correction

Processes biases decrease overall when the climatic data are corrected (in average 13 gC/m^2^/y for *AWBI*, 98 gC/m^2^/y for *GPP* and 34 gC/m^2^/y for respiration) and reach an error of 14% in average at plot level. Stressful years are listed in Appendix S11. The absolute value of the bias between simulation with SAFRAN and with uncorrected model is greater than the absolute bias with corrected models, for MPI and Hadgem models (respectively 72% and 68% in average any process combine) and CNRM models for *GPP* and *AWBI* 75% in average, meaning that climatic corrections decrease the bias to SAFRAN simulations (Fig 5). This result is more nuanced for CNRM model for *Resp* (only 50% of SER shows a bias improvement for respiration).

**Figure 5.**
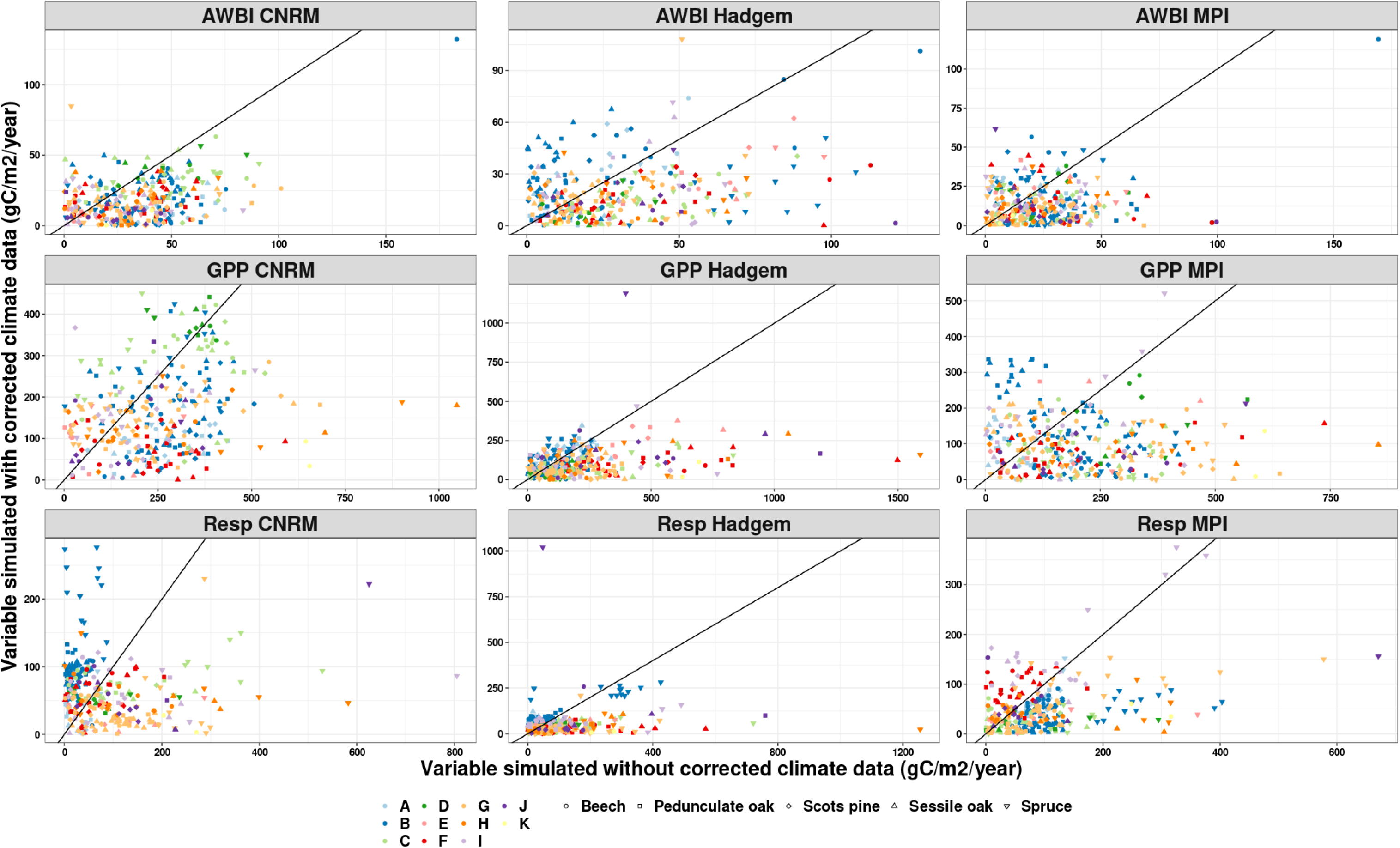
Comparison of process bias between simulation with uncorrected data and corrected data during four most stressful years, for each process and each model. Shapes represent species and GRECO are indicated by color. The black line is the line 1:1. Points below the line indicate a decreasing bias between uncorrected and corrected climate data. Points above the line indicate an increasing bias between uncorrected and corrected climate data.

##### Simulation bias and studies scales importance

Correction of modelling climate data divide almost by two percentage of process bias, for each scale and each process (Appendix S12). Bias percentage of process simulated with corrected modelling climate data differs between scales (plot, SER, GRECO) and process studied (Fig 4b). At plot level, bias of process are in average 10% for *GPP*, 8% for *Resp* and 23% for *AWBI* (Appendix S12), but can reach 90% for *AWBI* of monospecific oaks stand and *Resp* and *GPP* of monospecific pedunculate oak and beech stands. At regional scale (SER, GRECO and France), bias percentage are in average 9% for *GPP*, 8% for *Resp* and 19% for *AWBI* for SER and 10% for *GPP*, 8% for *Resp* and 20% for *AWBI* for GRECO and 10% for *GPP*, 8% for *Resp* and 18% for *AWBI* France. Standard error decreases when aggregation increases. Difference of bias percentage between scales varies depending on monospecific stands: from the highest bias percentage for sessile oak stand *AWBI* (from 31% for plot scales to 25% for regional scales) to lowest bias percentage for spruce stand *GPP* (from 4% for plot scales and for France scales). Percentage of *AWBI* bias are higher than percentage of *GPP* and *Resp* bias, at every scale.

##### Higher bias percentage compare to average year

Uncertainties are higher at every scale for stressful years compare to average year (Fig. 4-a,b and Appendix S12). Error can be twice as large for *AWBI* at plot scale or can be only slightly larger for *GPP* and *Resp* at regional scales (SER or GRECO).

## 4 Discussion

### 4.1 Scale and climatic stress intensity affect impact model outputs (biases)

Processes biases decrease overall when the climatic data are corrected and reach an error of 8% in average at plot level. In terms of percentage the impact model output errors after climate data correction are satisfactory low especially for *GPP* and respiration (respectively 6 and 7 %). *AWBI* prediction errors are slightly larger, with 11% in average.

This means that quantile-quantile and bias correction of climate data are essential and efficient to reduce biases in impact model predictions (De Cáceres et al., 2018; Ivanov & Kotlarski, 2017; Ruffault et al., 2014). Research on effective bias corrections of climate data is therefore a very important topic. These corrections methods should also be evaluated on different impact models.

#### 4.1.1 Assessing the effects of spatial scale

The question of the relationship between aggregation of models output at larger scale and biases is already present in the literature (Heuvelink & Pebesma, 1999 on soil process modelling). According to our results, larger scale means less large errors (outliers), but not a systematic decrease in bias average in general.

Note, however, that increasing the spatial scale and aggregation level allow to decrease the effect of climate data differences on *AWBI* (especially for sessile oak monospecific stands) and on *GPP* for monospecific beech stands. To our knowledge these findings are new and there is no study allowing to compare our results with others studies testing different levels of aggregations.

On one hand, in order to obtain less dispersion in CASTANEA predictions (outliers) aggregation at regional scale seems a good idea since outliers are very sensitive to local high climatic differences between SAFRAN and climatic modeling data. In fact aggregation decreases the statistical weight of extremes biased values. On the other hand monospecific stand areas per ecological regions are only available for a limited number of GRECOs (from 3 to 7 out of 11 GRECO, depending on the species) and a limited number of SER (from 4 to 10 out of 91 SER, depending on the species). However, species are present in practically all GRECOs: from 8 to 10 out of 11 GRECO, depending on the species. The aggregation method therefore allows us to study a representative part of French territory focusing on ecological regions where a computed forest area by species is available. .

In our work, we focused to emphasize bias percentage difference of fluxes (*GPP*, *AWBI*, respiration) between several spatial scales. This point is central, because forest manager questions require responses at different scales (as suggested in Bellassen et al. (2011) or in Bolte et al. (2009)): from the plot scale (management of a few hectares of forest), to the regional scale (supplying the wood industry). Understanding predictions quality at these different scales will make it possible to give answers in a different ways depending on the question asked. There are some limits to the approach used in this paper. For example, aggregation method involves regional data on monospecific forest area by species, yet the standard deviation on the area can be very large (Appendix S10), and the information or species may also be missing (see above). If we are only interested in comparisons, as is the case in this article, this is not a problem. But we must keep in mind that in absolute value predictions at the larger scale (regional level like SER or GRECO) adds de facto an additional error. In this respect, all considered, predictions with the least bias may not necessarily be the ones at largest scale.

#### 4.1.2 Focus on the driest years

In addition, we focused on the bias generated during driest years. Droughts are predicted to be more frequent (Dubrovský et al., 2014; Giorgi & Lionello, 2008; Polade et al., 2017), and it is therefore crucial to quantify models errors and to improve their prediction quality. Uncertainties comparison between stressful years and average years are very rare. For example, Jung et al (2007) highlighted that the major discrepancies are related, among other things, to the water stress effects in the most part of terrestrial ecosystem. According to our work, dry years have a lower prediction quality than average years at plot level: a higher error of 4 more percentage points on *GPP*, 1 on respiration and 12 on *AWBI*. This trend is in agreement with the results of Jung et al., (2007).

We observed that climate of the driest years of each climate dataset (Appendix S4) are mostly similar when *T* and *Rg* dynamics are considered but depends on the geographic area and climatic model for *Prec*, *V* and *RH*. The problem is mostly present for uncorrected climate data (for *V*) and in mountain area (for *Prec* and *RH*). In particular, in the uncorrected data, summer *RH* is largely underestimated in the majority of GRECOs (A-G) and winter *RH* is overestimated in GRECO H and I. The corrections decrease the differences between modelling climatic data and SAFRAN data, in regards to magnitude and seasonal variation, even if an overestimation still remains significant (p-value<0.0005 with Student test) in mountain ecological regions (Pyrenees and Alps). Overall, the most stressful years are equivalent between corrected modelling climate dataset and SAFRAN dataset. The large differences on *V*, *RH* and *Prec* in uncorrected climate data is larger and not limited to mountain areas. The larger bias of process predictions during stressful years may be due to large differences between modelling meteorological data and SAFRAN data.

#### 4.1.3 Correlation between climate data and impact model output biases

We observed that the SERs with the greatest errors due to climatic data biases are essentially in the Mediterranean rim and the South-East of Massif Central (Appendix S5-S6-S7). In order to assess the effect of climatic variables, we tried to evaluate if a correlation could be found between climatic variables (*T, RH, Rg, V, Prec*) and the process biases (*GPP, Resp, AWBI*). Methods and results are developed in Appendix S9.

Results show that correlation between climate and process bias are almost all significant (72/75), with an average of 0.18. Correlation are mostly positive (61/75), meaning larger biases in climatic models induces larger biases in impact model ouputs. The rare negative correlation (11/75) are mostly weak (−0.15 in average), except for Sessile oak and Scots pine stand *AWBI* and precipitation (correlation of −0.3). The strongest positive correlation is between biases on Scots pine, spruce, pedunculate and sessile oak stand respiration and mean annual temperature, and also Spruce stand *GPP* and temperature. It can be explained by the large effect of temperature on photosynthesis and respiration (Delpierre et al. 2012, Dusenge et al., 2019).

### 4.2 Process-based model prediction in context of global change

Process-based models take into account direct effect of climate on photosynthesis and respiration, contrary to a great number of models: for example only 9 to 49 models studied in Albrich et al. (2020) are considered as biogeochemical models, i.e. process-based models. For this reason, they represent promising prospects for understanding and predicting forest ecosystems functioning changes. Studies have already tried to predict trends of forest productivity (Madani et al., 2018; Prislan et al., 2019; Tei et al., 2017), carbon storage (Gustafson et al., 2017; Lindner et al., 2014; Lo et al., 2019) or C, H2O and N dynamics (Dong et al., 2019) in the decades to come.

Biases quantification on impact models outputs caused by the quality of the climate data used as input is rarely studied (Glotter et al., 2014; Palma et al., 2018; Stéfanon et al., 2015). Such studies are crucial, however, in order to be able to nuance interpretations of predictions, especially since the conclusions are contrasted. According to Stéfanon et al. (2015) goodness of fit of CASTANEA derived beech distribution, can be between 70% (when using meteorological data from the Aladin climate model) and less than 10% (when using meteorological data from the WRF climate model). Results are similar for the other model used (BIOMOD, a niche model). This means that the climate model used for prediction has a great importance (see also Glotter et al., 2014 for American yield). Palma et al. (2018) on the other hand has shown with 3-PG (Landsberg and Waring, 1997; Sands and Landsberg, 2002) that datasets of simulated climate can have minimal reduction of performance (in this study RACMO and WRF). Note that they used stock and not fluxes in their comparison, contrary to other cited studies.

It is also important to keep in mind that not all climate models allow the same quality of estimation. For example in our study the use of the CNRM climate model as input (with corrections) induced larger biases on the impact model simulations than the other climate models tested in our study (quite twice more than Hadgem for *GPP* and *Resp*). It is therefore important to compare impact model outputs used with climatic models input data against impact model outputs used with observed climatic data over an historical period. This allows to quantify the uncertainty associated with climatic data (see Jung et al., 2007) and is the prerequisite condition before making predictions about the future period.

More specifically, this work makes it possible to associate an estimate of biases on CASTANEA simulations when using modelling climate data on French territory and to quantify simulation errors for each climate model.

As explained in the introduction, uncertainties in impact model outputs are accumulated along a “cascade of uncertainty” (Lindner et al., 2014; Reyer, 2013), composed of uncertainty sources at each step of the chain. In this work, we focused only on one single step of this “cascade of uncertainty”: the effect of climatic input data biases on impact model output biases. As a matter of fact, for extreme climatic years (considering temperature and precipitation), bias of climate data may be the highest source of uncertainties (Iizumi et al., 2017). However, uncertainty of the impact model itself can also have a strong influence on the ouputs. Previous studies focused on CASTANEA uncertainties due to process-based model parameters (Dufrêne et al 2005).

Finally, we would like to remind the reader that we used only one single process-based model, CASTANEA, which limits the genericity of our results. In fact, inter-model comparisons studies show that the same set of climate data could induce different responses to the same question depending on the impact model used (see Jung et al 2007 or Cheaib et al 2012). It might be instructive to replicate our approach with other mechanistic models and more broadly with other impact models driven by climate model data.

## 5 Conclusion

Our work highlighted several crucial points in the development of process-based models predictions in the context of climate change. First, a simple correction of the climatic data allows obtaining a significant improvement in the predictions of the CASTANEA model (57 gC/ha/year in average, in 78% of the case). Second, a study at different scales showed that work on a regional scale (SER, GRECO, France) reduce biases variance due to climate model, but do not significantly reduce the average bias. Third, we were able to highlight that the bias on model’s predictions when using different climate models increases for the driest years compared to average years. These results are similar regardless of the process considered. Fourth, biases (in %) for wood growth (*AWBI*) are higher than those for *GPP* and respiration.

Therefore using impact model to make predictions using modelling climate as input is relevant and useful provided that the relevant corrections are performed. This study performed on an historical period was necessary to assess the relevance of the method (Jung et al., 2007) and prepare prediction studies over future periods (2020-2100).

## Authors contributions

MJ conceived the original question and simulation setup of this study. NMStP processed the modelling climate dataset. MJ, CF, NMStP and ED designed the research, developed the methodology; MJ processed and analysed the data; MJ, CF, ND, NMStP and ED led the writing of the manuscript. All authors contributed critically to the drafts and gave final approval for publication.

## Acknowledgements

This study strongly benefitted from the help of G. Marie. We greatly thank team EV of ESE research unit for helpful discussions. This study was funded by the project MOPROF (French Agence De l’Environnement et de la Maîtrise de l’Énergie, ADEME). We thank also T Audinot and JD Bontemps for their help in handling of IFN dataset.

