## Supplementary Materials for "« Idol with feet of clay »: reliable predictions of forest ecosystem functioning require flawless climate forcings"

#### Appendix S1 : CASTANEA calibration results

We performed a calibration of carbon allocation to aboveground biomass in CASTANEA for *Fagus sylvatica*, *Picea abies*, and *Pinus sylvestris*. We used IFN inventory, i.e. only even aged regular monospecific stand plots. Dataset is made up of individual 5-year tree rings increment inventories between 1970-2017. Then we divide in two the whole inventory for each species: one quarter for calibration and three quarters for validation, except for *Pinus sylvestris* where we used one-half for calibration and one-half for validation. To do this calibration we compared observed and simulated radial increment of mean tree, using function *optim* from package *stats*.

| Species | Calibration plots number | Validation plots number | RMSE calibration | RMSE validation |
|---|---|---|---|---|
| Fagus sylvatica | 1036 | 3109 | 0.16 | 0.19 |
| Pinus sylvestris | 2562 | 2563 | 0.18 | 0.17 |
| Picea abies | 752 | 2511 | 0.31 | 0.20 |

We computed RMSE, slope of linear model (simulated radial increment explained by observed radial increment) and $R^2$, for calibration and validation subpart of dataset.

#### Appendix S2 : GRECO and SER

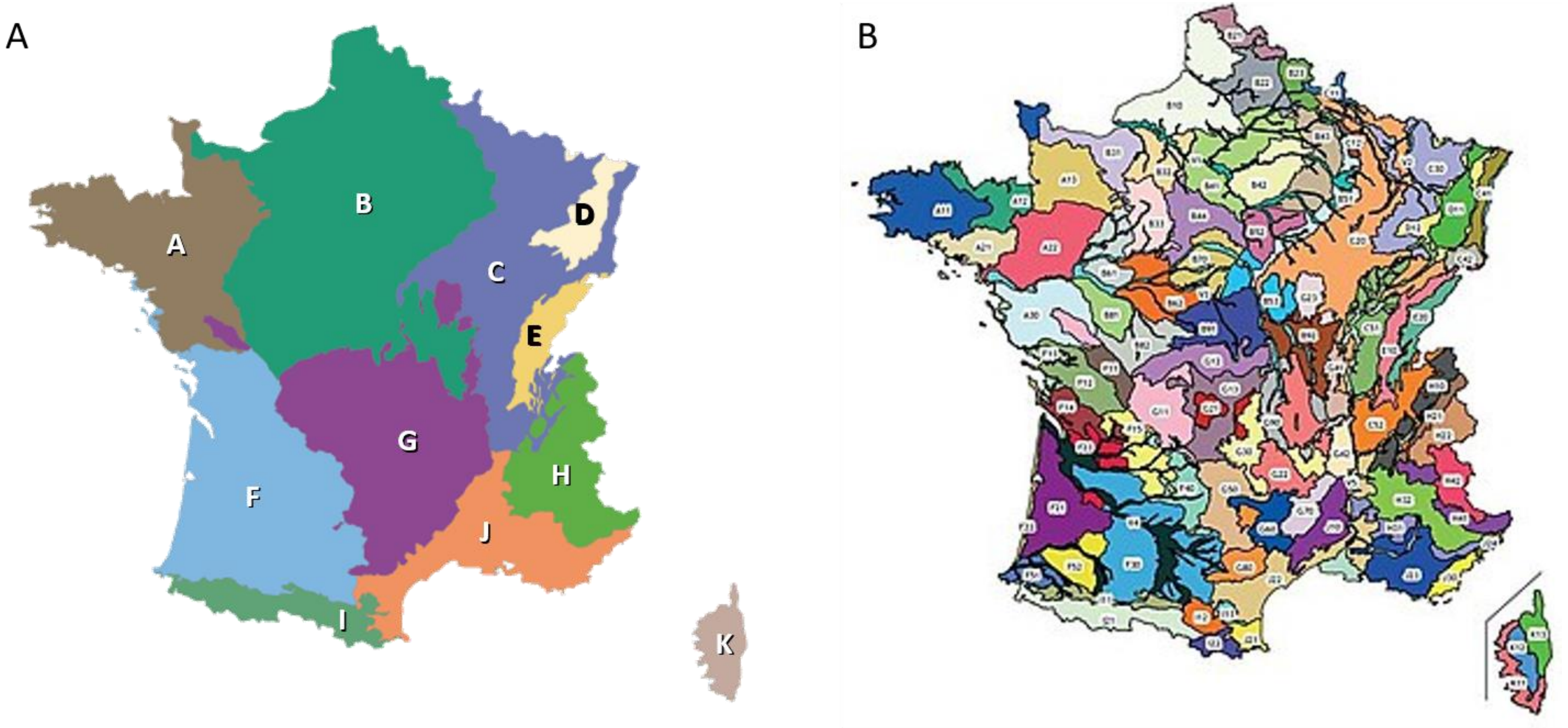


**GRECO** **SER**

*Maps of geographical delimitation of GRECO (Grande Region ECOlogique), macro-regional scale, (A) and SER ("Sylvo-Eco-Region" or Ecological Forest Region, meso-regional scale (B), from the website (*[https://inventaire-forestier.ign.fr/](https://inventaire-forestier.ign.fr/)*)*

### Appendix S3 : Computing details of equations 6 and 7

To study scale effect on quality of prediction (objective n°2, see above), we focused on two regional scales SER and GRECO, to understand aggregation effect on two scales. At the regional level, stand age distribution could have a strong effect on prediction because processes are strongly affected by stands age in CASTANEA and can interact with climate effects. For following analyses we considered four age classes (20-60, 60-120, 120-180, >180). We tested different age class, and results are similar.

We compute the proportion of bias compared to variable level at regional scale as in eq 3, using total surface of the species ($S_{sp}$), mean level mean level per hectare of variable V for all plots $i$ of a given age class ($mean_i(V)$) and proportion of this age class ($\%_{sp,s,age}$):

$$\delta_{mod,v,sp,s} = V_{mod,sp,s} - V_{saf,sp,s}$$

$$= S_{sp} * \sum_{age} \%_{sp,s,age} * mean_i(V_{mod,sp,s,age}) - S_{sp} * \sum_{age} \%_{sp,s,age} * mean_i(V_{saf,sp,s,age})$$

$$= S_{sp} * \sum_{age} \%_{sp,s,age} * \left(mean_i(V_{mod,sp,s,age}) - mean_i(V_{saf,sp,s,age})\right)$$

$$= S_{sp} * \sum_{age} \%_{sp,s,age} * mean_i(V_{mod,sp,s,age} - V_{saf,sp,s,age})$$

$$= S_{sp} * \sum_{age} \%_{sp,s,age} * mean_i(\delta_{mod,v,sp,s,age}) \text{ (a)}$$

Then,

$$p.biais_{mod,v,sp,s} = \delta_{mod,v,sp,s} / V_{saf,sp,s}$$

$$= (S_{sp} * \sum_{age} \%_{sp,s,age} * mean_i(\delta_{mod,v,sp,s,age})) / (S_{sp} * \sum_{age} \%_{sp,s,age} * mean_i(V_{saf,sp,s,age}))$$

$$= \sum_{age} \%_{sp,s,age} * mean_i(\delta_{mod,v,sp,s,age}) / \sum_{age} \%_{sp,s,age} * mean_i(V_{saf,sp,s,age}) \text{ (b)}$$

We need to assign a total area of monospecific area per region to each age class and each species. The monospecific stand area is not available by age class and species, but only by species. We therefore decided to compute the area by age class and by species using the total specific area and proportion of monospecific stand plot for each age class and species. Then we extracted total surface by *SER* and *GRECO* from IFN data (see Appendix S8). The ratio of plots number in each age class ($nb_{age}$) to the total plots number ($nb_{tot}$) is then determined for each type of monospecific stand by computing this proportion annually and averaging over the 10 years of inventory available (between 2006 and 2017). This gives, for each year $i$, SER $s$, age class *age* and species *sp*:

$$\%_{sp,s,age,i} = nb_{age,sp,s,i}/nb_{tot,sp,s,i} \text{ (c)}$$

The calculations are the same for GRECO level and France level.

#### Appendix S4: Seasonal variation of climatic variables during stressful year

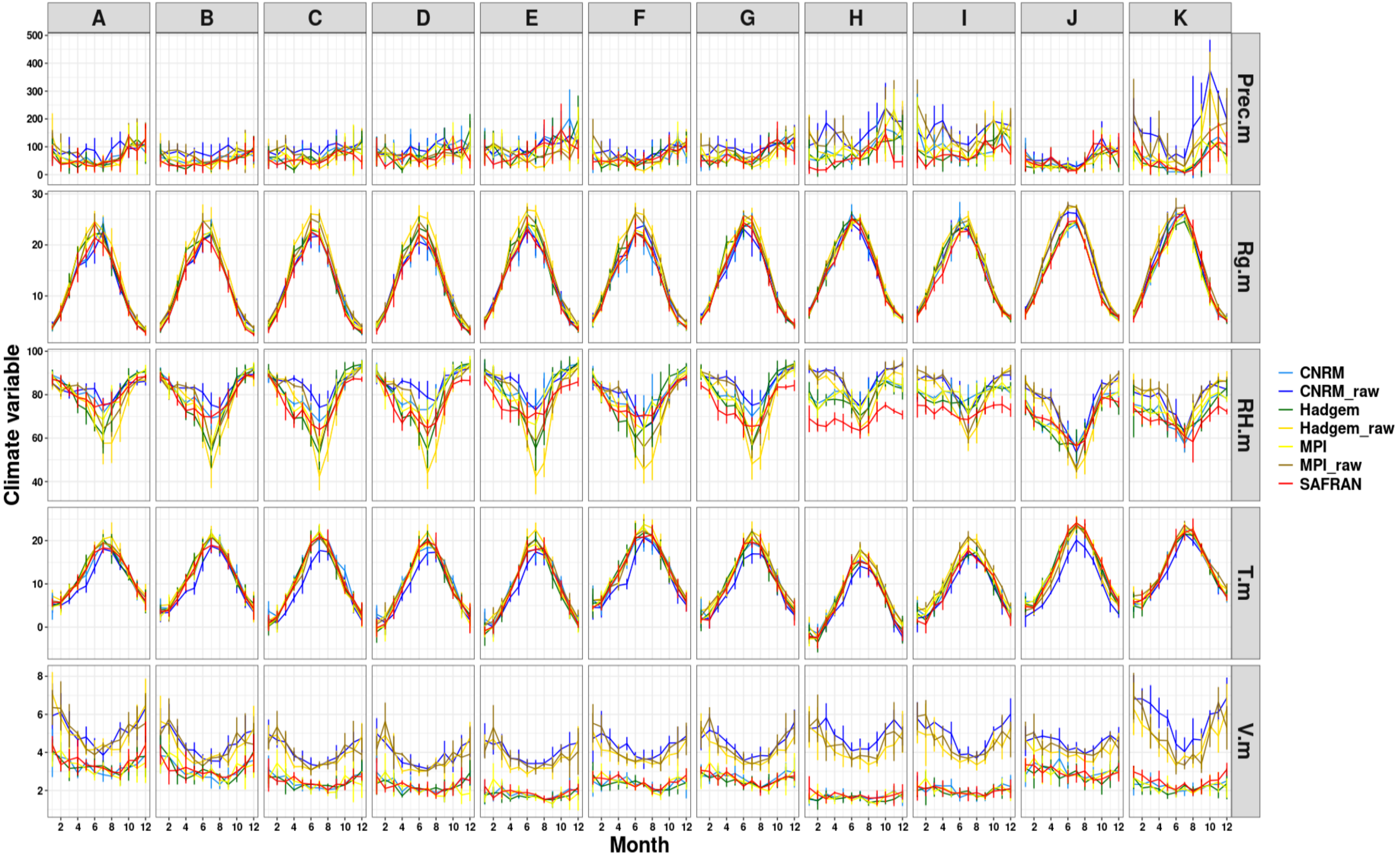


*Monthly climatic variables for radiation (m.Rg), relative humidity (m.RH), mean temperature (m.T) and Precipitation (Prec), for climatic model before and after correction and for each GRECO.*

### Appendix S5 : Maps of GPP bias

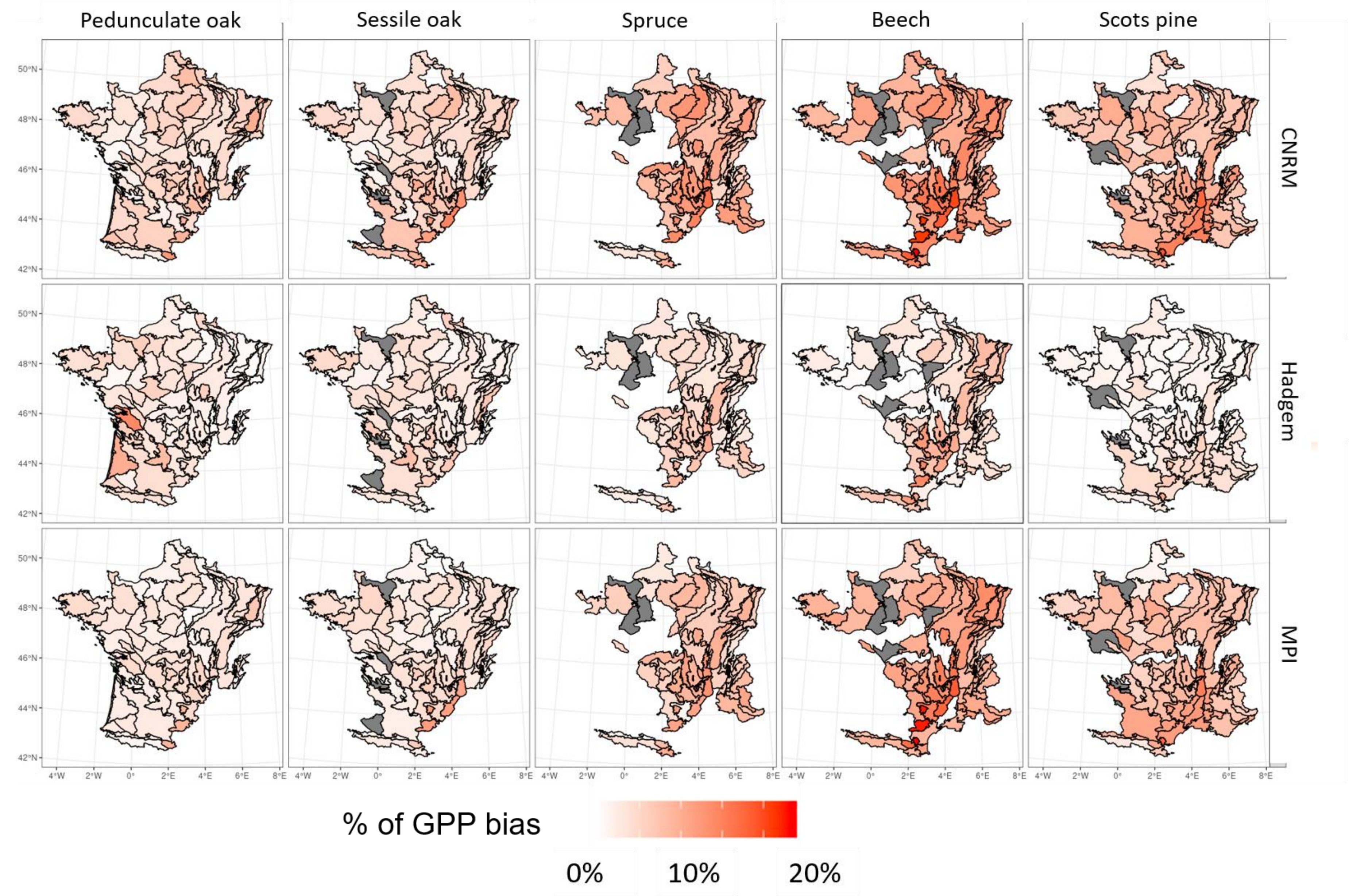


*Maps of average GPP bias percentage at plot scale, comparing simulations with climatic models and with SAFRAN data, for each model and each species. Polygon correspond at SER and color at uncertainties level.*

### Appendix S6 : Maps of respiration bias

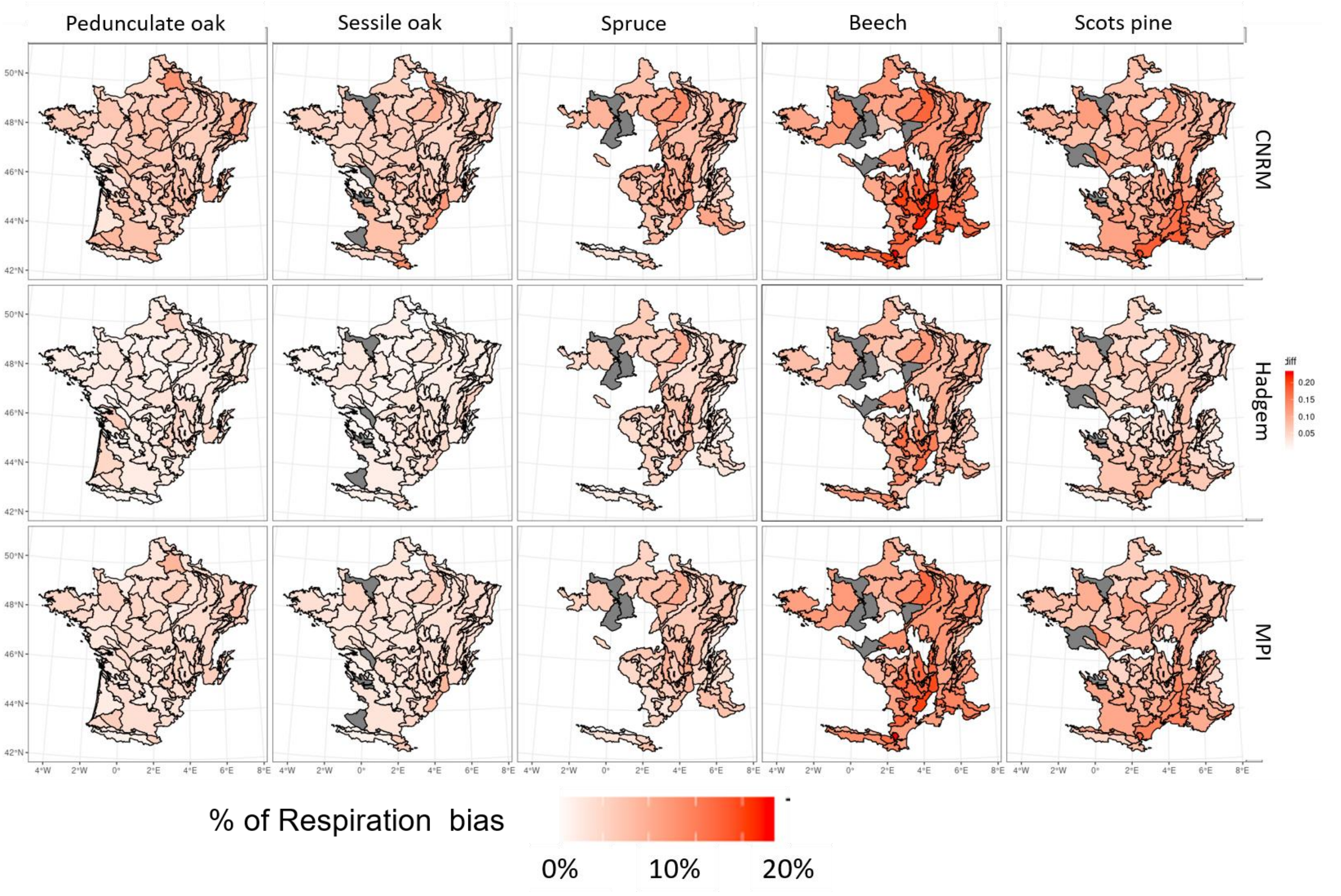


*Maps of average Respiration bias percentage at plot scale, comparing simulations with climatic models and with SAFRAN data, for each model and each species. Polygon correspond at SER and color at uncertainties level.*

### Appendix S7: Maps of AWBI bias

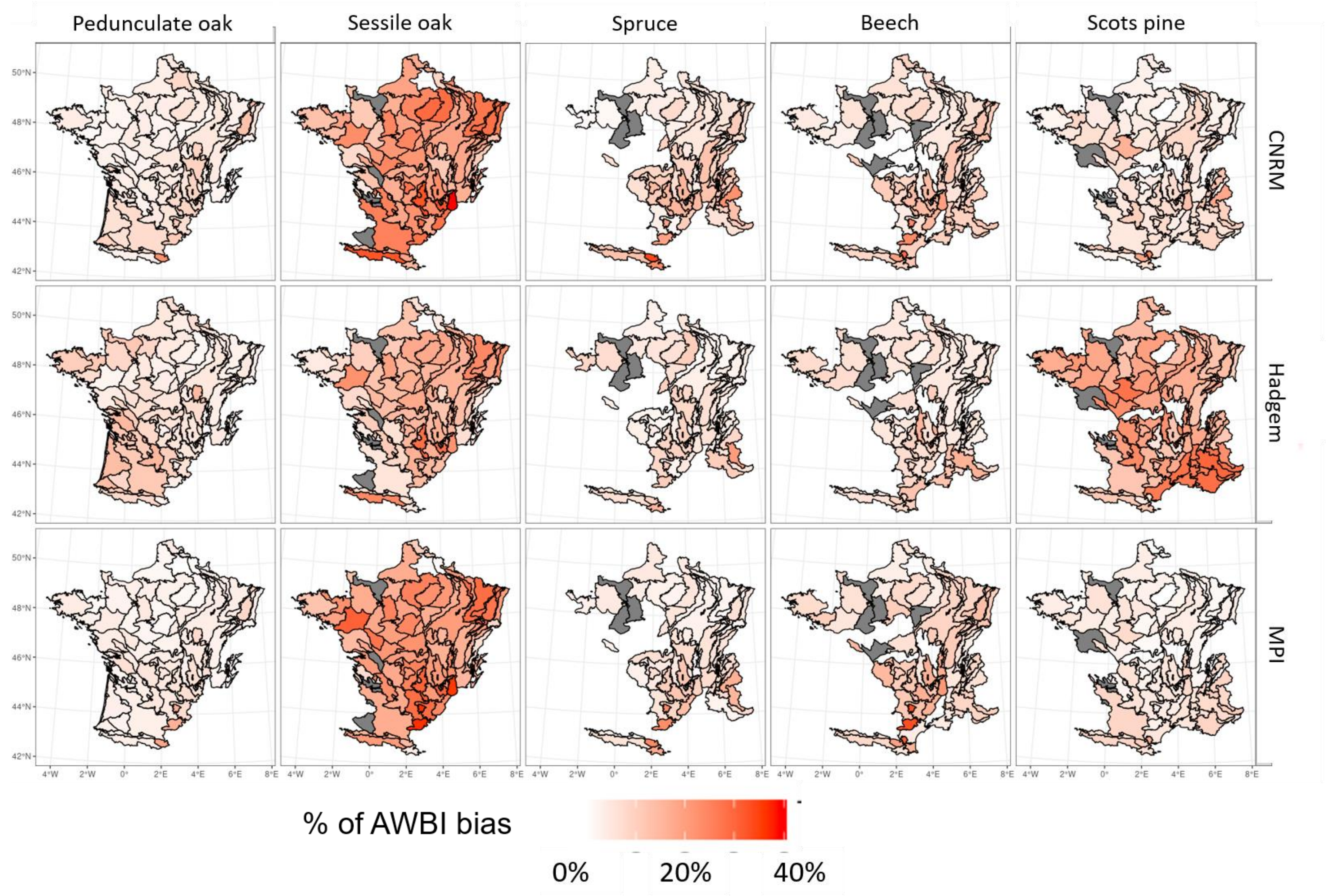


*Maps of average ABWI bias percentage at plot scale, comparing simulations with climatic models and with SAFRAN data, for each model and each species. Polygon correspond at SER and color at uncertainties level.*

### Appendix S8: Climate model and SAFRAN variables caracteristics

a)

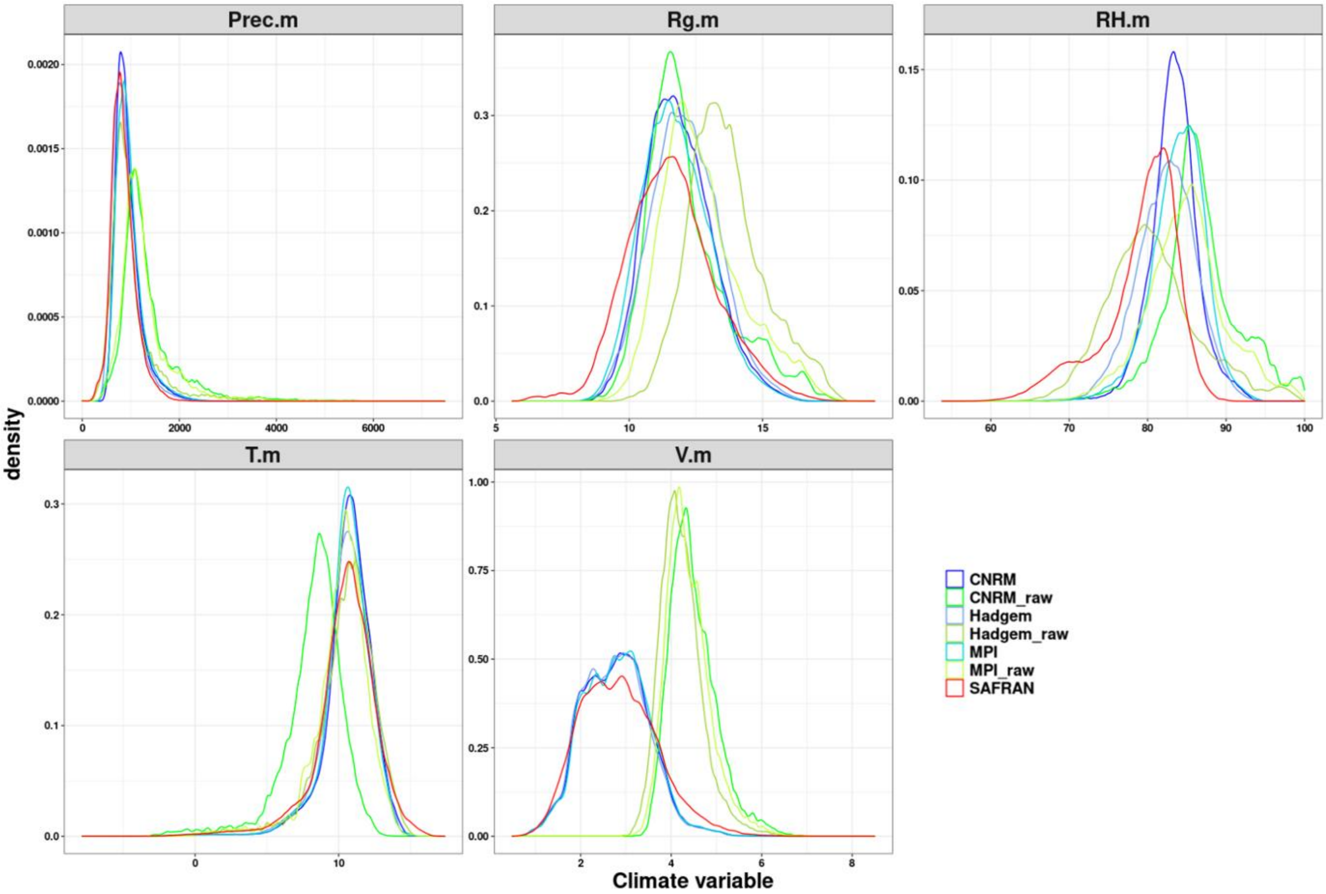


*Spatial distribution of climatic variables for radiation (Rg), relative humidity (RH), mean temperature (T) and Precipitation (Prec)*

b)

| GCM | From | RCM | Projected climatic changes for Europe |
|---|---|---|---|
| CNRM-CERFACS-CNRMCM5 | Centre National de recherche météorologiques | RCA4 | Moderate summer temperature increase, Slight summer precipitation increase |
| MOHC-HadGEM2-ES | Met Office Hadley Center | RCA4 | High summer temperature increase, High summer precipitation decrease |
| MPI-M-MPI-ESM-LR | Marx Planck Institute | RCA4 | High summer temperature increase, Medium summer precipitation decrease |

*Table represents chosen climate model characteristics (adapted from Fargeon et al 2020)*

### Appendix S9 : Correlation between climate and process bias

We decided to qualify relationship between process bias and climatic bias in SER with the highest percentage of bias (higher than 40%). First, climatic variables are averaged over period (1960-2010), by plot. Secondly, climatic bias is computed similarly to (1), for each variable (*V*), model (*mod*), species (*sp*) and each plot:

$$\delta_{mod,v,sp} = V_{mod,sp} - V_{saf,sp} \quad (1b)$$

We wanted to understand which climatic variables affect process percentage of bias (eq 4 in the main document) in function of climate bias (1b). Then we decided to quantify for each species and each process the correlation between process bias and climate bias.

| | Scots pine | | | Beech | | | Spruce | | | Pedunculate oak | | | Sessile oak | | |
|---|---|---|---|---|---|---|---|---|---|---|---|---|---|---|---|
| | GPP | Resp | AWBI | GPP | Resp | AWBI | GPP | Resp | AWBI | GPP | Resp | AWBI | GPP | Resp | AWBI |
| T | 0.2 | 0.35 | 0.07 | 0.2 | 0.35 | 0.07 | 0.2 | 0.35 | 0.07 | 0.2 | 0.35 | 0.07 | 0.2 | 0.35 | 0.07 |
| RH | 0.42 | 0.3 | -0.02 | 0.42 | 0.3 | -0.02 | 0.42 | 0.3 | -0.02 | 0.42 | 0.3 | -0.02 | 0.42 | 0.3 | -0.02 |
| Rg | 0.21 | 0.16 | 0.05 | 0.21 | 0.16 | 0.05 | 0.21 | 0.16 | 0.05 | 0.21 | 0.16 | 0.05 | 0.21 | 0.16 | 0.05 |
| V | 0.32 | 0.28 | 0.04 | 0.32 | 0.28 | 0.04 | 0.32 | 0.28 | 0.04 | 0.32 | 0.28 | 0.04 | 0.32 | 0.28 | 0.04 |
| Prec | 0.31 | 0.22 | -0.05 | 0.31 | 0.22 | -0.05 | 0.31 | 0.22 | -0.05 | 0.31 | 0.22 | -0.05 | 0.31 | 0.22 | -0.05 |

*Table represents correlation between climate bias and process bias at plot scale for each species and each process. Number represents every significant correlation and bold number correspond to strong positive correlation (>0.5) or negative correlation (<-0.5).*

### Appendix S10 : GRECO and SER areas by species

a)

| | A | B | C | D | E | F | G | H | I | Total |
|---|---|---|---|---|---|---|---|---|---|---|
| P. oak | 81 (±13) | 253 (±23) | 68 (±12) | - | - | 185 (±22) | 134 (±18) | - | - | 734 (±41) |
| S. oak | 52 (± 10) | 423 (±28) | 146 (±18) | 26 (±7) | - | 34 (±10) | 118 (±17) | - | - | 814 (±41) |
| B | - | 61 (±11) | 110 (±15) | 57 (±11) | 33 (±9) | - | 129 (±18) | 66 (±15) | 127 (±18) | 618 (±39) |
| S | - | - | 47 (±10) | 35 (±9) | 47 (±10) | - | 105 (±16) | 74 (±14) | - | 327 (±28) |
| SP | - | 75 (±13) | - | - | - | - | 132 (±19) | 220 (±27) | - | 503 (±38) |

b)

| | A11 | B10 | B32 | B33 | B41 | B52 | B53 | B62 | B70 | B91 | B92 | C20 | C30 | D11 | D12 | E20 | F21 | F30 | F52 | G11 | G13 | G22 | G30 | G60 | H10 | H30 | H41 | I21 |
|---|---|---|---|---|---|---|---|---|---|---|---|---|---|---|---|---|---|---|---|---|---|---|---|---|---|---|---|---|
| P. oak | 21(±6) | - | - | - | - | - | - | - | 68 (±12) | - | - | - | 23 (±7) | - | - | - | 38 (±11) | 33 (±10) | 32 (±9) | 33 (±9) | 32 (±8) | - | - | - | - | - | - | - |
| S. oak | - | - | 37 (±9) | 40 (±9) | 32 (±8) | 38 (±9) | 44 (±9) | 36 (±9) | 35 (±8) | 53 (±10) | 27 (±7) | 79 (±13) | 32 (±8) | - | - | - | - | - | - | - | - | - | - | - | - | - | - | - |
| B | - | 28 (±8) | - | - | 21 (±6) | - | - | - | - | - | - | 76 (±12) | - | 29 (±8) | 28 (±8) | - | - | - | - | - | - | 33 (±9) | 30 (±8) | - | - | - | - | 95 (±15) |
| S | - | - | - | - | - | - | - | - | - | - | - | - | - | 28 (±8) | - | 36 (±9) | - | - | - | - | - | 35 (±9) | - | - | 38 (±10) | - | - | - |
| SP | - | - | - | - | - | - | - | - | 40 (±9) | - | - | - | - | - | - | - | - | - | - | - | - | 42 (±11) | - | 41 (±11) | - | 157 (±23) | 41 (±12) | - |

*Tables represents surface (thousand hectares) of monospecific stand in each ecological each species (rows) and region (columns): GRECO (a) and SER (b)*

### Appendix S11 : Most stressful years

| model | A | B | C | D | E | F | G | H | I | J | K |
|---|---|---|---|---|---|---|---|---|---|---|---|
| NUM_SAF | 1976 | 1976 | 1976 | 1976 | 1976 | 2005 | 1976 | 2003 | 1967 | 2003 | 1961 |
| | 2011 | 2011 | 2003 | 2003 | 2003 | 1976 | 2003 | 2004 | 2005 | 2005 | 2017 |
| | 1959 | 1996 | 2015 | 1964 | 2004 | 2011 | 2015 | 1976 | 1986 | 2006 | 2003 |
| | 2005 | 1959 | 1964 | 2015 | 2009 | 2003 | 2005 | 2005 | 2006 | 2004 | 1973 |
| CNRM | 2001 | 2009 | 2001 | 2001 | 2001 | 2001 | 2009 | 2001 | 2001 | 2001 | 1997 |
| | 1970 | 2001 | 1964 | 1964 | 2009 | 2009 | 1964 | 1964 | 1964 | 1985 | 1985 |
| | 2009 | 1974 | 2009 | 2009 | 1964 | 1964 | 1998 | 1972 | 1997 | 1974 | 1998 |
| | 1985 | 1970 | 2005 | 2006 | 2006 | 1981 | 2001 | 2009 | 1999 | 1998 | 1970 |
| Hadgem | 1975 | 1975 | 1966 | 2002 | 2002 | 1966 | 1966 | 1966 | 1987 | 1966 | 1966 |
| | 1990 | 1966 | 2002 | 1966 | 1966 | 1987 | 2002 | 1987 | 1966 | 1987 | 2001 |
| | 1967 | 2011 | 1967 | 1975 | 1967 | 2002 | 2011 | 1967 | 2006 | 1967 | 1987 |
| | 1966 | 1986 | 1975 | 1967 | 1986 | 2011 | 1995 | 1995 | 1995 | 1990 | 1984 |
| MPI | 2005 | 2005 | 2000 | 2000 | 2006 | 2012 | 2012 | 1973 | 2007 | 2012 | 2007 |
| | 1970 | 2000 | 2005 | 1985 | 1970 | 1970 | 2000 | 1970 | 1999 | 1973 | 1995 |
| | 2012 | 1973 | 1973 | 1970 | 2000 | 1973 | 2006 | 2012 | 2000 | 2002 | 1973 |
| | 2000 | 1963 | 1970 | 2005 | 2005 | 2000 | 2005 | 2002 | 2012 | 1964 | 1978 |
| CNRM_brut | 1970 | 2009 | 2009 | 2001 | 2001 | 2001 | 2009 | 2001 | 1964 | 2001 | 1997 |
| | 2001 | 1970 | 2001 | 2009 | 2009 | 2009 | 1964 | 1964 | 2001 | 1985 | 1985 |
| | 1989 | 2001 | 1964 | 1989 | 2006 | 1970 | 1974 | 1972 | 1997 | 1964 | 1998 |
| | 2009 | 1974 | 2005 | 1964 | 1964 | 1974 | 2005 | 2009 | 1985 | 1972 | 1970 |
| Hadgem_brut | 1975 | 1975 | 1966 | 2002 | 2002 | 1966 | 1966 | 1966 | 1987 | 1966 | 1966 |
| | 1990 | 1966 | 2002 | 1966 | 1966 | 1987 | 2002 | 1987 | 1966 | 1987 | 1982 |
| | 1966 | 2000 | 1975 | 1975 | 1986 | 2002 | 1987 | 1983 | 2006 | 2011 | 1987 |
| | 1987 | 2011 | 1967 | 1967 | 1967 | 2011 | 2011 | 2011 | 1995 | 1982 | 1984 |
| MPI_brut | 2005 | 2005 | 2000 | 2000 | 2006 | 2012 | 2012 | 1973 | 2000 | 2012 | 2007 |
| | 1970 | 1973 | 2005 | 2005 | 1970 | 2000 | 2000 | 2012 | 2007 | 1973 | 2012 |
| | 2012 | 2000 | 1970 | 1970 | 2000 | 1970 | 2006 | 1985 | 1999 | 2007 | 1973 |
| | 2000 | 1970 | 1973 | 2006 | 2005 | 1979 | 1973 | 1970 | 2012 | 2002 | 2000 |

*Table represents the four driest year of periode 1960-2010 for each climate data source and each GRECO*

### Appendix S12 : Table of mean bias percentage

| | | Average years | | | Stressful years | | |
|---|---|---|---|---|---|---|---|
| **scale** | **correction type** | **$NPP_{wood}$** | **GPP** | **Resp** | **$NPP_{wood}$** | **GPP** | **Resp** |
| **plot** | **Corrected** | 11 (±12) | 6 (±4) | 7 (±5) | 23 (±34) | 10 (±8) | 8 (±7) |
| | **Non-corrected** | 25 (±21) | 13 (±11) | 12 (±11) | 43 (±53) | 18 (±15) | 13 (±12) |
| **SER** | **Corrected** | 9 (±7) | 6 (±4) | 7 (±4) | 19 (±13) | 9 (±6) | 8 (±6) |
| | **Non-corrected** | 23 (±16) | 13 (±9) | 11 (±9) | 36 (±24) | 17 (±11) | 13 (±9) |
| **GRECO** | **Corrected** | 9 (±5) | 6 (±3) | 7 (±4) | 20 (±12) | 10 (±7) | 8 (±5) |
| | **Non-corrected** | 23(±10) | 14 (±8) | 13 (±7) | 36(±15) | 18 (±9) | 13 (±7) |
| **France** | **Corrected** | 9(±5) | 6(±3) | 7(±4) | 18(±6) | 10(±4) | 8(±4) |
| | **Non-corrected** | 22(±6) | 13(±2) | 12(±3) | 35(±9) | 17(±4) | 13(±3) |

*Table of mean bias percentage (and standard deviation) of simulation compare to process levels simulated with SAFRAN, by process (column), scale and correction type (row), for average years and most stressful year*
